# The generation of a Bcl11a lineage tracing mouse model

**DOI:** 10.1101/2021.07.26.453822

**Authors:** Sara Pensa, Pentao Liu, Walid T. Khaled

## Abstract

The transcription factor B-cell lymphoma/leukaemia 11A (BCL11A) has essential functions in physiological processes as well as haematological and solid malignancies, however, its contribution to tissue development and tumour progression is poorly understood. Here we show the generation of a *Bcl11a* lineage tracing mouse model to allow for *in vivo* tracking of *Bcl11a*-expressing cells and their progeny. We validate the model in the mammary gland by using flow cytometry and whole-tissue 3D imaging to locate labelled cells after induction of tracing in early development. We show that *Bcl11a* is predominantly expressed in long-lived luminal progenitors which populate mammary alveoli upon pregnancy, confirming *bona fide* labelling of *Bcl11a* cells. The *Bcl11a* lineage tracing mouse model therefore provides a powerful resource to study *Bcl11a* cells in development, homeostasis, and cancer.

## Introduction

The transcription factor B-cell lymphoma/leukaemia 11A (BCL11A) is a zinc finger protein mainly expressed in the brain and hematopoietic cell lineages ^1,2^. BCL11A is part of the ATP-dependent chromatin remodelling mammalian SWI/SNF complex ^3^, which enables nucleosome restructuring, ensuring DNA is accessible for transcription replication and DNA repair ^4^.

*BCL11A* is essential for B-cell development and plays a crucial role in the foetal to adult haemoglobin switch^5–7^. Evidence is also accumulating of its role in various other physiological processes, including a developmental role in the skin and in determining dendritic cell fate ^8,9^. It is also implicated in the 2p15-p16.1 microdeletion syndrome, characterised by intellectual disability, Type II diabetes and β-hemoglobinopathies, particularly sickle cell disease (SCD)^1^. Targeting *BCL11a* has been implemented to regulate foetal haemoglobin (HbF) levels in erythrocytes in a transgenic humanized sickle cell mouse model and in SCD patients, thus providing preliminary evidence of the therapeutic effects of such a strategy for β-hemoglobinopathies ^10,11^.

In addition, *BCL11A* is highly expressed in some haematological malignancies and malignant solid tumours and is associated with poor clinical prognosis^1^.We have previously shown that *BCL11A* is a triple negative breast cancer (TNBC) gene^12^. *BCL11A* is overexpressed in TNBC including basal-like breast cancer (BLBC), and its genomic locus is amplified in up to 38% of BLBC. *BCL11A* overexpression in cell lines promotes tumour formation whereas its knockdown supresses tumour formation in xenograft models. Importantly, *Bcl11a* deletion severely decreases tumour formation in a mouse model of TNBC^12^.

We have also characterised *BCL11A* role as a lung squamous carcinoma (LUSC) oncogene^13^. We showed that *BCL11A* is upregulated in LUSC but not in lung adenocarcinomas and that knockdown of *BCL11A* in LUSC cell lines abolishes xenograft tumour growth and its overexpression *in vivo* leads to lung airway hyperplasia and the development of reserve cell hyperplastic lesions^13^. At the molecular level we found *BCL11A* to be a target of SOX2, a key driver of LUSC^14^, and we showed that it was required for the oncogenic role of SOX2 in LUSC^13^. Furthermore, we showed that BCL11A and SOX2 interact at the protein level and that together they co-regulate the expression of several transcription factors, including *STED8*, and inhibition of STED8 selectively affects LUSC growth^13^.

Despite its implication in various pathophysiologies, *BCL11A* contribution to tissue development and tumour initiation is poorly characterised. Lineage tracing is the most widely used technique to track stem and progenitor cells and their progeny *in vivo* and to explore the contribution of these cells to development, homeostasis, and cancer^15^. The fates of individual cells can be traced using both spatial and temporal control. Well characterised gene promoters drive expression in the cell types of interest, while inducible systems allow for temporal control. The main inducible systems used for stem cell biology are a tamoxifen-regulatable version of the *Cre* recombinase that is engineered behind an endogenous or transgenic promoter and a tetracycline-responsive reverse transactivator (rtTA) linked to a cell-specific promoter, incorporating a separate *TetO-Cre* allele.

Considering *BCL11A* important functions in tissue physiology, as well as its widespread implication in malignancies, we sought to generate a lineage tracing mouse model implementing the tamoxifen-inducible *Cre* as a strategy to trace *Bcl11a*-expressing cells fate, and validated the labelling strategy in mammary epithelial cells.

## Results

The *Bcl11a* lineage tracing mouse model was generated by knocking-in an *IRES-CreERT2* cassette into the 3` UTR region of *Bcl11a*, which we hereafter refer to as *Bcl11a*^*CreERT2*^ (Figure 1A). The *Bcl11a*^*CreERT2*^ allele was generated by introducing an IRES-CreERT2-polyA-FRT-PGK-E7-Neo-polyA-FRT cassette into the 3′UTR of *Bcl11a* locus. Gene targeting was carried out in AB2.2 cells (129 background). Correct targeting was confirmed by long-range PCR and targeted clones were used for blastocyst injections. Chimeric mice were then crossed to wildtype mice (C57/B6) and germline transmission was confirmed (Figure 1B). These mice allowed us to perform lineage tracing when crossed to the *Rosa26-LSL-tdTomato* reporter mice (*tdTom*) to generate double transgenic mice (Figure 1C). Tamoxifen mediated CreERT2 activation induced tdTomato expression in *Bcl11a* expressing cells and any of their future progeny. We have shown before that *Bcl11a* is expressed predominantly in luminal progenitor cells and, in smaller proportion, in basal cells and differentiated luminal cells in the adult mammary gland^12^. To confirm that tamoxifen administration leads to true labelling of *Bcl11a* expressing cells, we checked If this was the case in the mammary gland by inducing lineage tracing in the adult with 3 injections on 3 consecutive days of tamoxifen (1 mg per injection). Mammary tissues were collected at 1 day (1 day p.i.) or 5 days (5 days p.i.) post injection and RNA was extracted from epithelial cells. At 1 day p.i. tdTomato^pos^ cells express higher levels of *Bcl11a* than tdTomato^neg^ cells (Figure 1D), indicating specific labelling. At this time point we would expect labelling of progenitor as well as differentiated, *Bcl11a*-expressing cells in the mammary gland, as previously reported^12^. At 5 days p.i., *Bcl11a* expression in tdTomato^pos^ cells is reduced compared to 1 day p.i., reflecting the persistence of the progenitors cells labelling after recycling of the mammary gland and loss of the *Bcl11a*-expressing differentiated cells after 5 days p.i. (Figure 1D).

**Figure 1.**
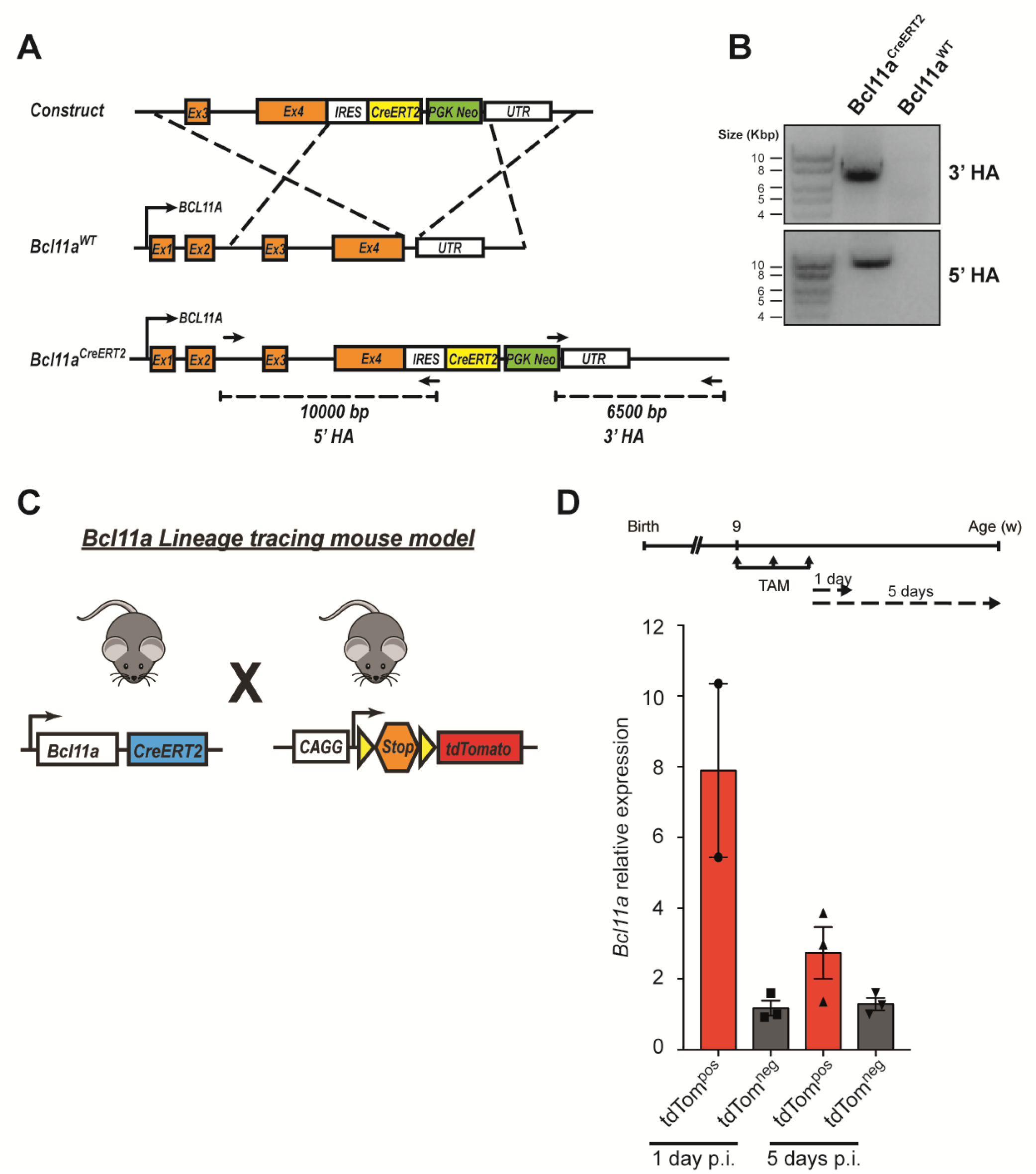
Generation and characterisation of Bcl11a^CreERT2^ mice by lineage tracing. **A**, schematic representation of the genetic strategy to insert the CreERT2 cassette into the 3’ UTR of the *Bcl11a* locus. **B**, long range PCR detecting the 5’ and 3’ homology arms (5’ HA and 3’ HA, respectively) of tissue samples from mice with the successful integration of the targeting cassette. The primers used in the long-range PCR are represented as arrows in the schematic in **A. C**, Schematic describing *Bcl11a* lineage tracing model. **D**, To induce *Bcl11a* tracing, mice were injected intraperitoneally with 3 low doses of tamoxifen on 3 consecutive days post puberty and tissues collected at 1 day (1 day p.i.) or 5 days (5 days p.i.) post injection. Epithelial cells were FACS sorted based on EpCAM expression and RNA extracted. The graph shows the levels of *Bcl11a* expression normalised to *Gapdh* in tdTomato^pos^ compared to tdTomato^neg^ cells from the same sample, as detected by qRT-PCR. Data are represented as mean ± SD of 3 technical repeats.

To confirm the identity of labelled cells, labelling was induced at puberty or in the adult with either 1 single intraperitoneal injection or 3 consecutive days of tamoxifen (1 mg per injection). Mammary tissues were collected at 1 day p.i. and fluorescence-activated cell sorting (FACS) analysis of the mammary compartments was performed (Figure 2A). One day post-labelling revealed that the vast majority (95% ± 1.6) of tdTomato^pos^ cells reside within the luminal compartment (Figure 2B, C) and are primarily luminal progenitors (Figure 2D). Interestingly, the contribution of tdTomato^pos^ cells to the luminal compartment increased 8 weeks after labelling (Figure 2E), further confirming the progenitor identity of the labelled *Bcl11a* luminal cells. To further assess the progenitor nature of *Bcl11a*-expressing cells, females were mated 8 weeks after tamoxifen injection and tissues were harvested at gestation day 14.5. We found that the number of tdTomato^pos^ luminal cells expanded significantly to constitute approximately 30% of the luminal compartment, showing that *Bcl11a*-expressing progenitor cells contribute to the alveologenesis process (Figure 2F).

**Figure 2.**
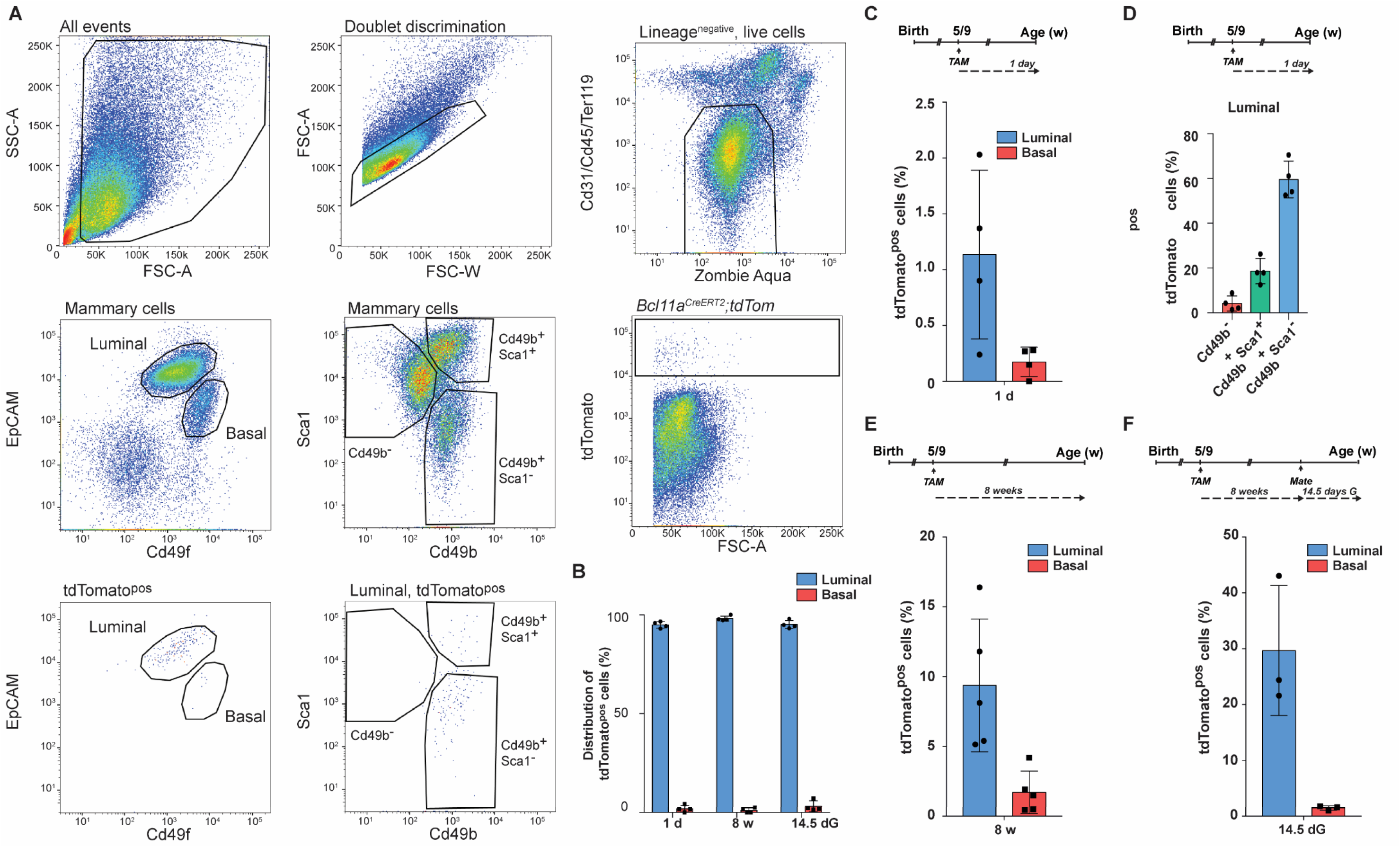
Bcl11a labels long-lived luminal progenitors that expand upon gestation. **A**, Representative plots showing the gating strategy used to select live, lineage negative, single tdTomato^pos^ cells in the mouse luminal and basal mammary epithelium, based on EpCAM and Cd49f staining of single cell preparations from lymph node divested mammary glands. FSC-W: forward scatter width, FSC-A: forward scatter area, SSC-A: side scatter area. Luminal differentiated and progenitor cells were identified based on Cd49b and Sca1 staining (Cd49b^-^, differentiated; Cd49b^+^ Sca1^+^ and Cd49b^+^ Sca1^-^, luminal progenitors). **B**, *Bcl11a* labelling was induced by intraperitoneal injection with 1 or 3 low doses of tamoxifen during or post puberty and tissues were collected at 1 day (1 d) and 8 weeks (8 w) post labelling and 14.5 days gestation (14.5 dG). The graph shows the distribution of tdTomato^pos^ cells within the epithelial compartment, analysed by FACS. Data are shown as mean percentage ± SD. **C**, to induce *Bcl11a* tracing, mice were injected as in **B** and tissues were collected 1 day post labelling. The graph shows the percentage of tdTomato^pos^ cells in the luminal and basal compartment as determined by FACS. Data are represented as mean ± SD. **D**, the graph shows the percentage of tdTomato^pos^ cells in differentiated (Cd49b^-^) or progenitor (Cd49b^+^, Sca1^+^ or Sca1^-^) luminal cells as determined by FACS 1 day post labelling. **E**, The graph shows the percentage of tdTomato^pos^ cells in the luminal and basal compartment as determined by FACS 8 weeks (8 w) post labelling. Data are represented as mean ± SD. **F**, *Bcl11a* labelling was induced and mammary glands were collected at 14.5 days gestation (14.5 dG). The graph shows the percentage of cells in the luminal and basal compartment, as determined by FACS. Data are represented as mean ± SD.

To gain insights into the spatial distribution of the labelled cells *in situ*, we performed optical tissue clearing of whole mammary glands followed by multicolour confocal 3D imaging^16,17^. By visualising the epithelium using EpCAM and cytokeratin-5 (K5), we observed the presence of individual cells and small clusters of tdTomato^pos^ cells throughout the luminal layer of the mammary gland (Figure 3A). To confirm *Bcl11a-*labelled cells contribution to the alveologenesis process we analysed the mice at 14.5 days gestation, where we could observe clusters of red cells in the mammary alveoli (Figure 3B).

**Figure 3.**
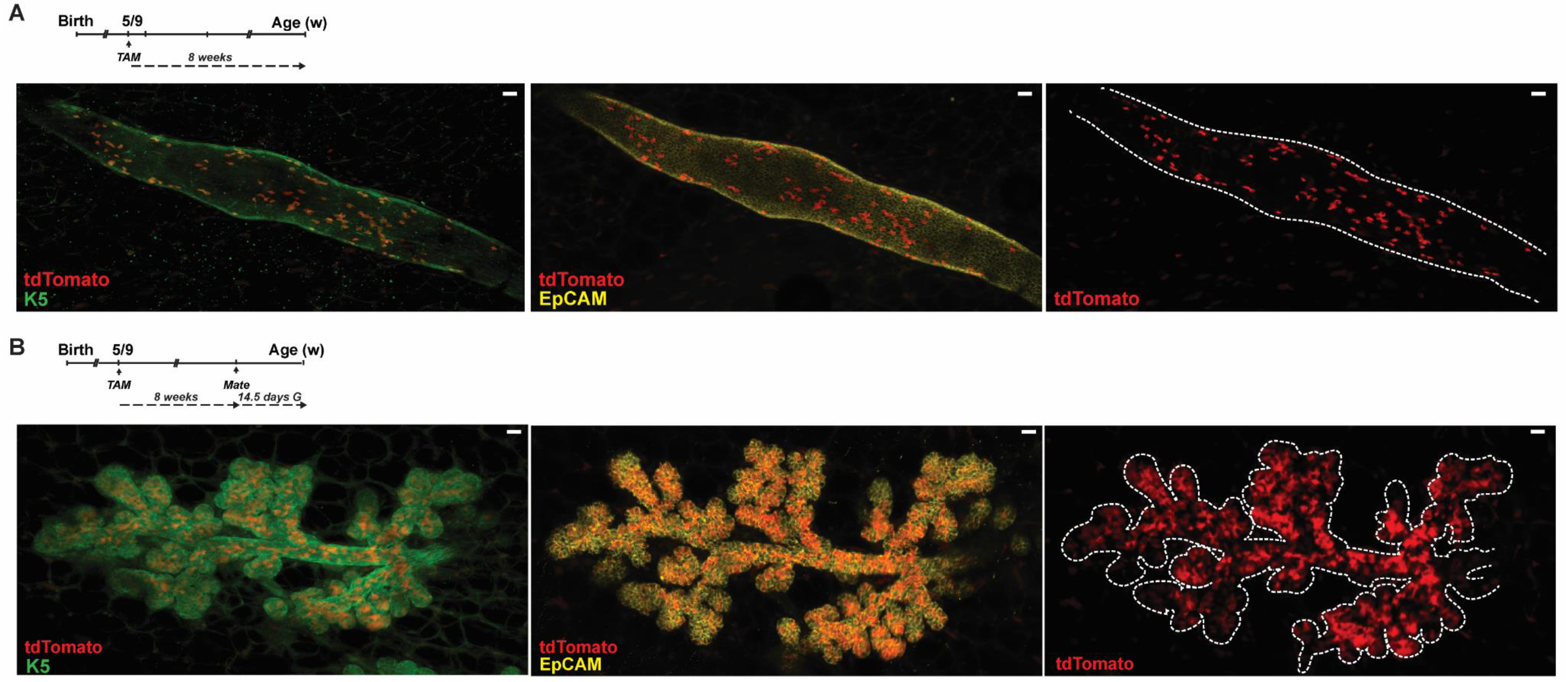
In situ localization of Bcl11a labelled cells in the mammary gland. **A**, *Bcl11a* tracing was induced as in **Figure 2** and tissues were collected at 8 weeks (8 w) post labelling. Confocal 3D imaging of tdTomato^pos^ cells of CUBIC-cleared whole glands immunostained with K5 and EpCAM are shown as overview; scale bars, 100 µm. **B**, *Bcl11a* labelling was induced and mammary glands were collected at 14.5 days gestation (14.5 dG). Confocal 3D imaging of tdTomato^pos^ cells of CUBIC-cleared whole glands immunostained with K5 and EpCAM are shown as overview; scale bars, 100 µm.

## Conclusion

Here we report the generation of a *Bcl11a* lineage tracing mouse model. This novel mouse model adds an important tool to study the biology of *Bcl11a* in health and disease. We validated this new lineage tracing mouse model by assessing labelling in mammary epithelial cells. The reporter mouse recapitulated previous results which report the expression of *Bcl11a* in luminal progenitor cells^12^. In addition, our analysis suggest that *Bcl11a* labelled cells are long-lived and contribute to pregnancy induced alveolar differentiation. The implication of this finding on the role of Bcl11a in tumour initiation remains to be explored. In conclusion, this new mouse model provides a powerful tool to explore the role of *Bcl11a* in normal development and cancer in the multiple tissues it has been proposed to play a key role.

## Author contribution

S.P. conceptualised and performed all the experiments. The *Bcl11a*^*CreERT2*^ mice were made by W.T.K. whilst in P.L.’s laboratory. S.P. and W.T.K. wrote the manuscript with input from the other authors. W.T.K. conceptualised and supervised the study.

## Acknowledgements

We would like to thank the staff at Sanger Institute, Research Service Facility (RSF) and the staff at the Cambridge NIHR BRC Cell Phenotyping Hub for their constant support and assistance. W.T.K. is funded by a CRUK Career Establishment Award (C47525/A17348), CRUK Small Molecule Drug Discovery Project Award (C47525/A25850), Breast Cancer Now Project Grant (2017MayPR907), University of Cambridge and Magdalene College, Cambridge.

## Materials and Methods

### Mouse experiments

All experimental animal work was performed in accordance to the Animals (Scientific Procedures) Act 1986, UK and approved by the Ethics Committee at the Sanger Institute. Correct targeting was confirmed by long-range PCR. All the primers used for genotyping are listed in Table S1. Details of the lineage tracing experiments are summarised in Table S2. The *Rosa26-LSL-tdTomato* mouse line was obtained from the Jackson Laboratory (JAX 007905) (Ref 25?). For each experiment, the animals used were heterozygous for *Bcl11a*^*CreERT2*^ and homozygous for *Rosa26-LSL-tdTomato*. Lineage tracing was induced at puberty (5 weeks of age) or in the adult (9 weeks of age) with either 1 single intraperitoneal injection or 3 injections on 3 consecutive days of tamoxifen (1 mg per injection) in corn oil. F or each time point *Bcl11a*^*CreERT2*^ wildtype control littermates were injected at the same time as experimental animals and tissues collected to set background levels of tdTomato fluorescence both in FACS and whole tissue IF experiments. All the primers used for genotyping are listed in Table S1. All mice were housed in individually ventilated cages under a 12:12 h light–dark cycle, with water and food available ad libitum and euthanized by terminal anaesthesia.

### Mammary gland dissociation into single-cell suspension

Lymph node divested mammary glands (excluding the cervical pair) were dissected from the mice and digested O/N in DMEM/F12 (Gibco) + 10 mM HEPES (Gibco) + 1 mg ml^-1^ collagenase (Roche) + 100 U ml^−1^ hyaluronidase (Sigma) (CH) + gentamicin (Gibco) at 37 °C. After the lysis of red blood cells in NH_4_Cl, cells were briefly digested with warm 0.05% Trypsin-EDTA (Gibco), 5 mg ml^−1^ dispase (Sigma) and 1 mg ml^−1^ DNase (Sigma) and filtered through a cell strainer (BD Biosciences).

### Cell labelling followed by flow cytometry and sorting

Single-cell suspensions were incubated in HF medium (Hank’s balanced salt solution (Gibco) + 1% fetal bovine serum, Gibco) + 10% normal rat serum (Sigma) for 20 min on ice to pre-block before antibody staining. All antibody incubations were performed for 10 min on ice in HF media. Mouse mammary cells were stained with the following primary antibodies: Cd31-biotin (eBioscience, clone 390, 1 µg ml^−1^, 1:500); Cd45-biotin (eBioscience, clone 30F11, 1 µg ml^−1^, 1:500); Ter119-biotin (eBioscience, clone Ter119, 1 µg ml^−1^, 1:500), EpCAM-APC/Cy7 (Biolegend, clone G8.8, 0.5 µg ml^−1^, 1:500), Cd49f-BV421 (Biolegend 313623, 2 µg ml^−1^, 1:100), Cd49b-AF488 (Biolegend, clone HMα2, 1 µg ml^−1^, 1:500) and Sca1-AF647 (Biolegend, clone D7, 1 µg ml^−1^, 1:500). Cells were then stained with streptavidin-APC or Streptavidin-PE/Cy7 (BD-Biosciences, 0.4 µg ml^−1^, 1:500). 7-AAD (Sigma, 10 µg ml^−1^, 1:100) or Zombie Aqua (Biolegend, 1:100) were used to detect dead cells. Cells were filtered through a cell strainer (Partec) before sorting. Sorting of cells was done using a SH800Z sorter (SONY) after sorting calibration was performed with automatic setup beads (SONY) immediately prior to sorting, FACS analysis was performed on a FACS Aria Fusion. Single-stained control cells were used to perform compensation manually. Unstained cells and control animals were used to set gates. The gating strategies is reported in **Figure 2A**. Cells were sorted for RNA analysis or analysed for tdTomato expression in mammary luminal or basal compartments. FlowJo was used to analyse FACS data.

### Optical tissue clearing and wholemount immunostaining

Mammary tissue was dissected and cut into large pieces (∼15 × 15 × 2 mm) for immunostaining and clearing. CUBIC-based tissue clearing ^18^ was performed as previously described ^16,17^ and combined with wholemount immunolabelling for visualization of tdTomato cells. Briefly, CUBIC Reagent 1A was prepared using urea (Sigma, 10% (w/w)), N,N,N’,N’-tetrakis(2-hydroxypropyl)ethylenediamine (Sigma, 5% (w/w)), triton-X100 (VWR, 10% (w/w)) and NaCl (Sigma, 25 mM) in distilled water. CUBIC Reagent 2 was prepared using sucrose (Fisher Scientific, 44% w/w), urea (Sigma, 22% w/w), 2,2′,2″-nitrilotriethanol (Sigma, 9% w/w) and Triton X-100 (VWR, 0.1% w/w) in distilled water. Tissues were immersed in CUBIC Reagent 1A at 37 °C for 2-3 days. For immunostaining, samples were washed and subsequently blocked in PBS containing triton-X100 (0.5% (w/v)) and goat serum (10% (v/v)) overnight at 4 °C. Primary antibodies were diluted in blocking buffer at 4 °C for 4 days with gentle rocking. Tissue was washed (3 × 1 h) and incubated with Alexa Fluor conjugated secondary antibodies for 2 days, washed in PBS and transferred to CUBIC Reagent 2 at 37 °C for at least 1 day for refractive index matching. Samples were immersed in CUBIC Reagent 2 for imaging and were imaged within 2 weeks. The following primary antibodies were used for immunostaining: rat anti-EpCAM-AF647 (Biolegend, clone G8.8, 1 µg ml^−1^, 1:250); rabbit anti-K5 (Covance, PRB160P, 1:100). The following Alexa Fluor-conjugated secondary antibodies were purchased from Thermo Fisher Scientific and used 1:500: goat anti-mouse 488 (A11001); goat anti-rat 647 (A21247).

### Confocal microscopy and image analysis

Images of wholemount mammary glands were acquired using a Leica TCS SP8 and SP5 inverted confocal microscopes with a 40 × /1.3 HC PL APO objective lens. Laser power, line averaging and step increment were adjusted manually to give optimal fluorescence intensity for each fluorophore with minimal photobleaching. Image overview and 3D reconstructions were generated using ImageJ software.

### Preparation of RNA

Sorted cells were spun down and resuspended in RLT, and RNA was extracted using the RNeasy mini kit (for mouse cells) according to manufacturer instructions. DNA was degraded by adding 20U Rnase-free DnaseI (Roche) for 30 min at room temperature. DnaseI treatment was performed on columns.

### Preparation of cDNA and qRT-PCR

Total RNA was diluted to a final volume of 11 µl. 2 µl of random primers (Promega) were added after which the mixture was incubated at 65 °C for 5 min. A master mix containing Transcriptor Reverse Transcriptase (Roche), Reverse Transcriptase buffer, 2 mM dNTP mix and RNasin Ribonuclease Inhibitors (Promega). This mixture was incubated at 25 °C for 10 min, then 42 °C for 40 min and finally 70 °C for 10 min. The resulting cDNA was then diluted 1:2.5 in H_2_O for subsequent use. qPCR was performed using a Step-One Plus Real-Time PCR System (Thermofisher Scientific). Taqman (ThermoFisher Scientific) probes with GoTaq Real Time qPCR Master Mix (Promega) were used. The enrichment was normalised with control mRNA levels of *GAPDH* and relative mRNA levels were calculated using the ΔΔCt method comparing to control group. For the list of primers and probes see Table S1.

---

**Table S1.**
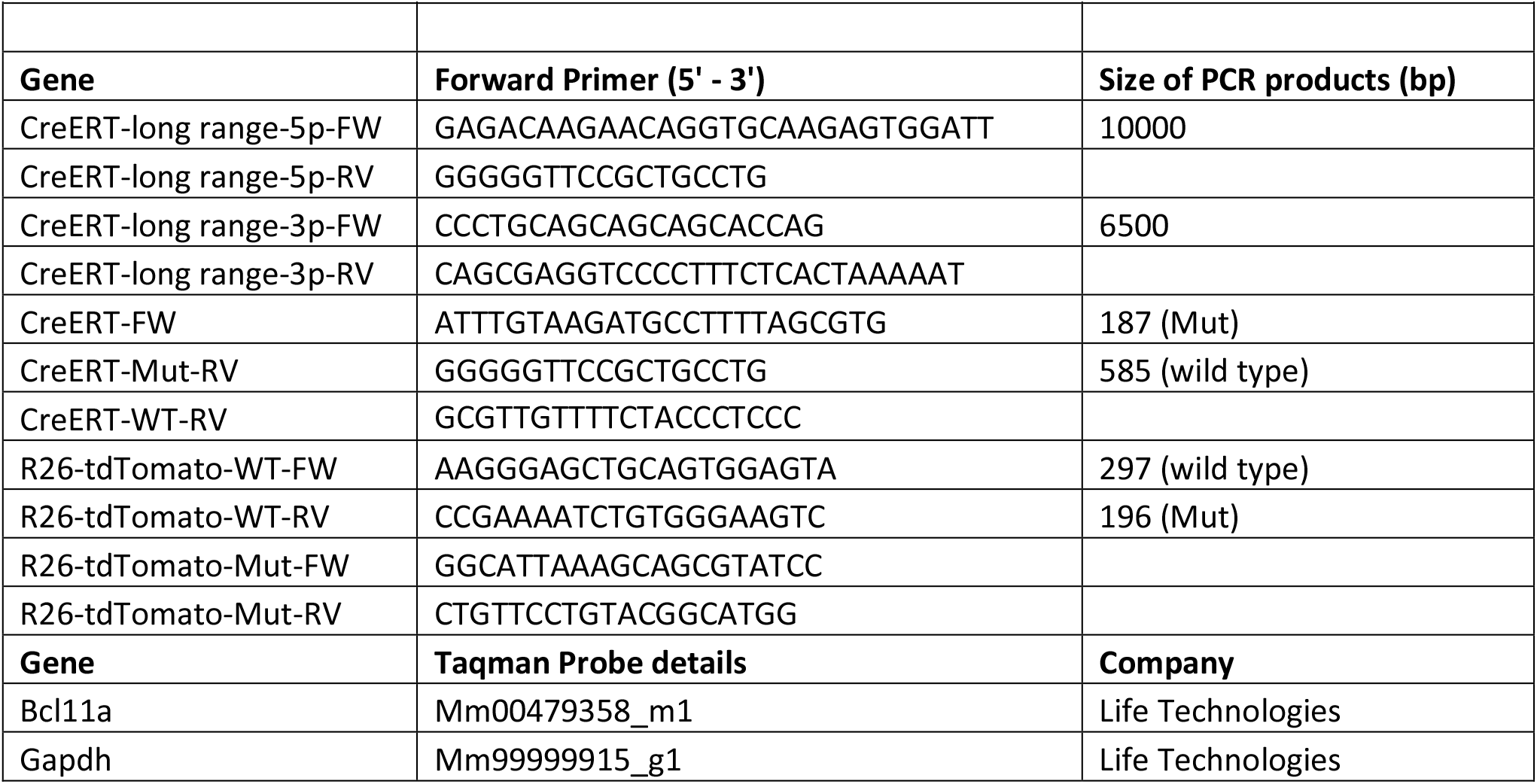
Genotyping primers list and qRT-PCR probes list.

**Table S2.** Lineage tracing data.

| Table S2 - Lineage tracing data |  |
| --- | --- |
| Chase group, 1 TAM injection | n of animals |
| 1 day | 4 |
| 5 days | 3 |
| Chase group, 3 TAM injection | n of animals |
| 1 day | 1 |
| 5 days | 1 |
| 8 weeks | 5 |
| 14.5 days gestation | 3 |

## References

1. Yin, J., Xie, X., Ye, Y., Wang, L. & Che, F. BCL11A: a potential diagnostic biomarker and therapeutic target in human diseases. Biosci. Rep. 39, BSR20190604 (2019).

2. Nakamura, T. et al. Evi9 encodes a novel zinc finger protein that physically interacts with BCL6, a known human B-cell proto-oncogene product. Mol. Cell. Biol. 20, 3178–3186 (2000).

3. Masliah-Planchon, J., Bièche, I., Guinebretière, J.-M., Bourdeaut, F. & Delattre, O. SWI/SNF Chromatin Remodeling and Human Malignancies. Annu. Rev. Pathol. Mech. Dis. 10, 145–171 (2015).

4. Tang, L., Nogales, E. & Ciferri, C. Structure and Function of SWI/SNF Chromatin Remodeling Complexes and Mechanistic Implications for Transcription. Prog. Biophys. Mol. Biol. 102, 122–128 (2010).

5. Liu, P. et al. Bcl11a is essential for normal lymphoid development. Nat. Immunol. 4, 525–532 (2003).

6. Yu, Y. et al. Bcl11a is essential for lymphoid development and negatively regulates p53. J. Exp. Med. 209, 2467–2483 (2012).

7. Sankaran, V. G. et al. Developmental and species-divergent globin switching are driven by BCL11A. Nature 460, 1093–1097 (2009).

8. Li, S. et al. Transcription Factor CTIP1/ BCL11A Regulates Epidermal Differentiation and Lipid Metabolism During Skin Development. Sci. Rep. 7, 13427 (2017).

9. Ippolito, G. C. et al. Dendritic cell fate is determined by BCL11A. Proc. Natl. Acad. Sci. U. S. A. 111, (2014).

10. Xu, J. et al. Correction of Sickle Cell Disease in Adult Mice by Interference with Fetal Hemoglobin Silencing. Science (80-.). 334, 993LP–996 (2011).

11. Esrick, E. B. et al. Post-Transcriptional Genetic Silencing of BCL11A to Treat Sickle Cell Disease. N. Engl. J. Med. 384, 205–215 (2021).

12. Khaled, W. T. et al. ARTICLE BCL11A is a triple-negative breast cancer gene with critical functions in stem and progenitor cells. Nat. Commun. 6, (2015).

13. Lazarus, K. A. et al. BCL11A interacts with SOX2 to control the expression of epigenetic regulators in lung squamous carcinoma. Nat. Commun. 9, 1–11 (2018).

14. Bass, A. J. et al. SOX2 is an amplified lineage-survival oncogene in lung and esophageal squamous cell carcinomas. Nat Genet 41, 1238–1242 (2009).

15. Kretzschmar, K. & Watt, F. M. Lineage tracing. Cell 148, 33–45 (2012).

16. Lloyd-Lewis, B. et al. Imaging the mammary gland and mammary tumours in 3D: optical tissue clearing and immunofluorescence methods. Breast Cancer Res. 18, 127 (2016).

17. Davis, F. M. et al. Single-cell lineage tracing in the mammary gland reveals stochastic clonal dispersion of stem/progenitor cell progeny. Nat Commun 7, 13053 (2016).

18. Susaki, E. A. et al. Whole-brain imaging with single-cell resolution using chemical cocktails and computational analysis. Cell (2014) doi:10.1016/j.cell.2014.03.042.

